# Identification of regulatory elements in primary sensory neurons involved in neuropathic pain

**DOI:** 10.1101/2022.09.09.507328

**Authors:** Kimberly E. Stephens, Cedric Moore, David A. Vinson, Bryan E. White, Zachary Renfro, Weiqiang Zhou, Zhicheng Ji, Hongkai Ji, Heng Zhu, Yun Guan, Sean D. Taverna

## Abstract

Chronic pain is a significant public health issue that is often refractory to existing therapies. Here we use a multiomic approach to identify cis-regulatory elements that show differential chromatin accessibility, and reveal transcription factor (TF) binding motifs with functional regulation in the dorsal root ganglion (DRG), which contain cell bodies of primary sensory neurons, after nerve injury. We integrated RNA-seq to understand how differential chromatin accessibility after nerve injury may influence gene expression. Using TF protein arrays and chromatin immunoprecipitation-qPCR, we confirmed C/EBPγ binding to a differentially accessible sequence and used RNA-seq to identify processes in which C/EBPγ plays an important role. Our findings offer insights into TF motifs that are associated with chronic pain. These data show how interactions between chromatin landscapes and TF expression patterns may work together to determine gene expression programs in DRG neurons after nerve injury.

## Introduction

Chronic pain remains a significant public health problem that directly affects over 20% of the US population and costs over $500 billion annually in health care and lost productivity (Yong et al., 2022, Institute of Medicine (U.S.). Committee on Advancing Pain Research Care and Education., 2011). Once established, chronic pain is often refractory to existing treatments and associated with alterations in mood and sleep patterns, poorer perceived health, and decreased quality of life (Pitcher et al., 2019, Gureje et al., 1998, Smith et al., 2001).

Nerve injury-induced chronic pain is characterized by complex activity-dependent plasticity and heightened excitability of neurons in the pain pathway. Context-dependent regulation of enhancer and/or repressive gene expression requires coordinated transcription factor binding to *cis*-regulatory elements (CREs). However, the transcriptional regulatory landscape which orchestrates these changes in gene expression in the context of nerve injury is just starting to be addressed (Stephens et al., 2021).

Epigenetic mechanisms are well-established regulators of a wide variety of physiological and pathological processes (Andersson et al., 2014). One major pathway of epigenetic modulation is the targeted addition or removal of small chemical post-translational modifications (PTMs) on individual nucleosomes (Strahl and Allis, 2000, Jenuwein and Allis, 2001). These histone PTMs can help alter the positioning of individual nucleosomes and therefore, facilitate access to CREs. In particular, mono-methylation of the lysine residue at position 4 on histone H3 (e.g., H3K4me1) is enriched at regulatory loci, and facilitates recruitment of the cohesin complex and other remodeling machinery (Heintzman et al., 2007, Local et al., 2018). While some pioneer transcription factors can recognize and bind to CREs within nucleosomal DNA, for non-pioneer transcription factors to recognize and bind to CREs, the target sequence must be unobscured by nucleosomes (Zovkic, 2021). Therefore, potential active regulatory regions may be identified by assays that identify accessible chromatin, such as the Assay for Transposase-Accessible Chromatin using Sequencing (ATAC-seq) (Buenrostro et al., 2013, Kwasnieski et al., 2014).

Primary sensory neurons in the dorsal root ganglion (DRG) play important roles in the development and maintenance of neuropathic pain. However, CREs, which likely mediate long-term changes in gene expression, have not been identified genome-wide in the DRG after nerve injury. In a well-established rat model of neuropathic pain induced by chronic constriction injury (CCI) of the sciatic nerve, we used ChIP-seq to identify H3K4me1 enrichment and ATAC-seq to comprehensively map chromatin accessibility at CREs in the lumbar DRGs, and compared findings to that in the naïve rats. We further integrated RNA-seq profiles to understand how differential chromatin accessibility after physical nerve injury may influence gene expression.We then performed motif analysis of the differentially accessible H3K4me1 enriched regions to identify enrichment of transcription factor binding motifs. Importantly, using a transcription factor protein array and ChIP-qPCR, we confirmed C/EBPγ binding to a differentially accessible sequence. Finally, RNA-seq was used to identify biological processes in which C/EBPγ plays an important role. These data provide valuable resources for our understanding and the further investigation of injury-induced changes in the regulatory landscape in primary sensory neurons.

## Results

### Genome-wide identification of chromatin accessibility in naïve and injured DRGs

Adult (i.e., 8-10 week-old) female Sprague Dawley rats underwent CCI surgery or were left unperturbed (i.e., naïve control). At 14 days following surgery, the ipsilateral lumbar (L4-L6) DRGs were removed from both groups. To identify regions of chromatin accessibility, we first performed ATAC-seq on the DRGs from naive rats and CCI rats (Figure 1). Two (naïve group) or three (CCI group) biological replicates were processed for a total of five ATAC-seq libraries. These libraries were sequenced to an average depth of 50.6 million total reads and generated 32 million unique reads that aligned to the rat genome (Supplemental table 1). We visualized the extent of similarity/dissimilarity of chromatin accessibility of the individual samples using the first two principal components from the principal component analysis of all genes (Supplemental figure 1A). The first two principal components accounted for approximately 62% of the total variance among the samples and produced distinct clusters of the samples by treatment group (i.e., naïve vs. CCI). We used MACS2 to call peaks that represent genomic regions of chromatin accessibility in each sample. A total of 118,329 unique regions of chromatin accessibility were identified in the CCI rats and 123,738 in the naive group. To ensure the stringency of our analysis, we only considered reproducible regions (Methods; Supplemental figure 1B). A total of 62,854 unique genomic regions of chromatin accessibility were identified across both naïve and CCI rats. Consistent with known enrichment profiles of active regulatory elements, the distance between chromatin accessible regions and TSS of the nearest gene suggests that these regions are concentrated in CREs (i.e., introns, intergenic regions) (Supplemental figure 1C, D). A smaller proportion of accessible regions was found in promoters and TSS of annotated genes. However, the ATAC-seq peaks near the TSS were of greater intensity than non-TSS peaks, which supports previous studies that find greater accessibility of chromatin around the TSS than in surrounding genomic regions (Thurman et al., 2012).

**Figure 1.**
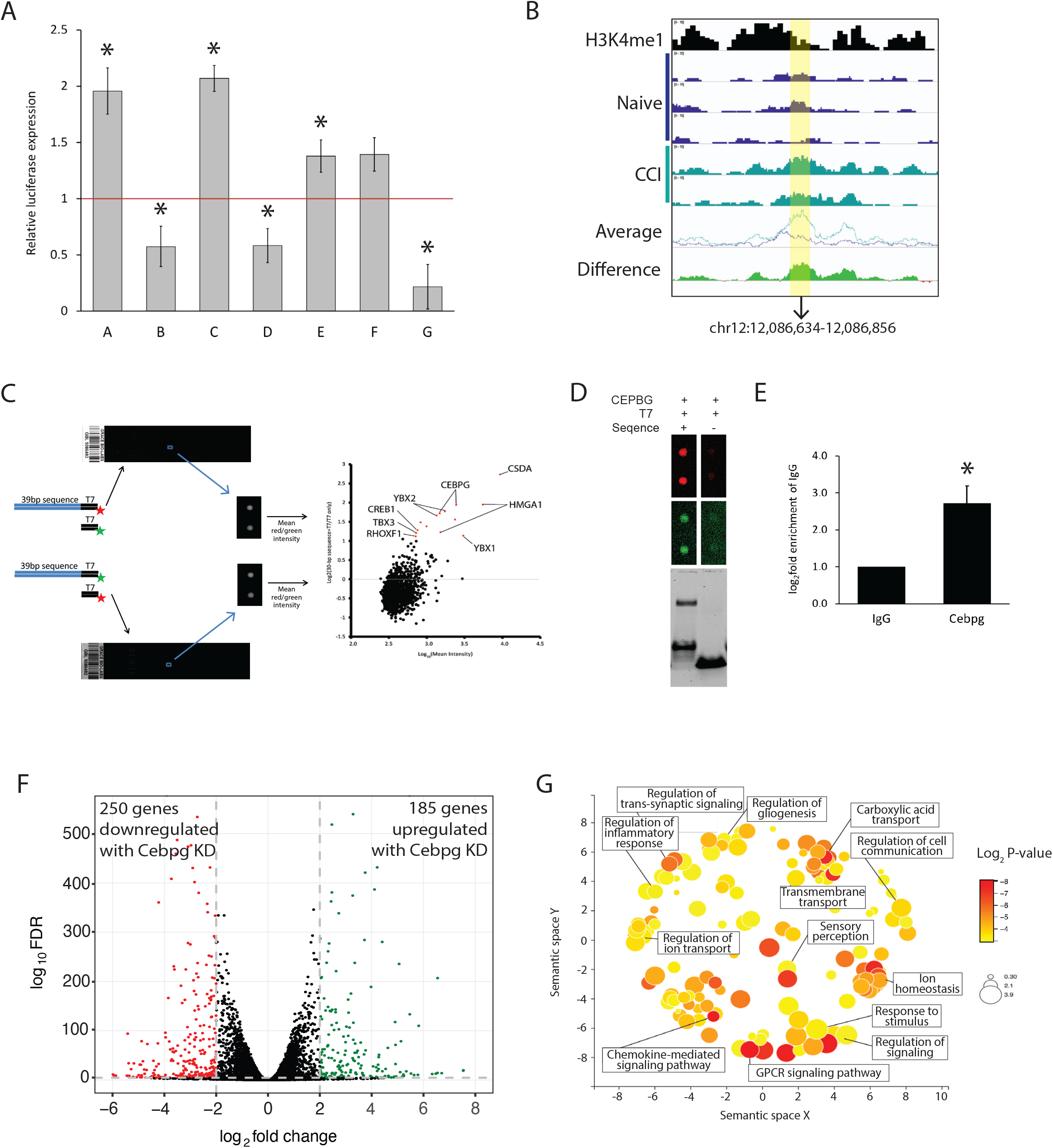
Chromatin accessibility in the rat DRG. Schematic of the experimental approach.

### Differential accessibility at gene promoters is associated with nociceptive processes

Analyses of our ATAC-seq on the DRGs from naive and CCI rats identified a union of 6,809 unique regions of open chromatin in annotated gene promoters between the groups (Figure 2A). Most (96.7%) of these promoter regions were accessible in both the CCI and naive groups (Figure 2A). The promoter regions of 108 genes contained open chromatin that was CCI-group specific, whereas 117 were naïve group-specific. Gene ontology (GO) analysis revealed that the 108 genes with CCI-specific accessibility were enriched for pain- and sensory-related processes (Figure 2A). Genes associated with the top 5 biological processes (as ranked by the *P* values) have well-established roles in persistent pain (e.g., the sodium voltage-gated channel Scn11a, and the transient receptor potential cation channels Trpv1 and Trpa1) (Figure 2B). For example, DRGs from CCI rats showed higher accessibility at the *Scn11a* promoter than the DRGs from naïve rats (left panel, Figure 2C), and this accessibility was associated with the higher gene expression (right panel; Figure 2C).

**Figure 2.**
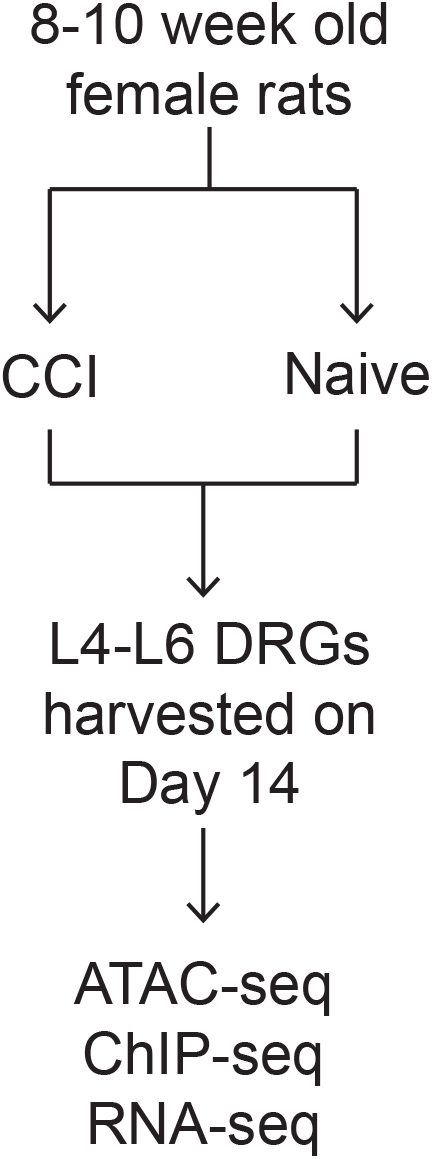
Changes in chromatin accessibility at gene promoters. A) Venn diagram shows the overlap of Naive and CCI accessible regions at annotated gene promoters (top). Bar plot showing the top 10 enriched Gene Ontology Biological Processes identified from 108 peaks associated only with annotated promoters in DRGs from CCI rats (bottom). The threshold for statistical significance was set to p-value < 0.01 (black horizontal line). B) Heatmap that shows the normalized accessibility and hierarchical clustering of the gene promoters that were enriched in the top 5 GO terms from (A). C) Normalized chromatin accessibility tracks at the Scn11a gene promoter (chr8:128,519,864-128,522,250). The individual and averaged ATAC-seq signal of the normalized bigwig files for each sample as displayed from the Integrated Genomics Viewer (left). The “difference” track was created by subtracting the average track from the naïve group from the average track of the CCI group. Green indicates an increase in accessibility in the CCI group compared to Naïve. The region highlighted in yellow (chr8:128,521,066-128,521,432) indicates the differentially accessible region between the Naïve and CCI samples. A box plot of the normalized, log_2_ transformed gene expression of Scn11a for each sample is provided on the right. D) Heatmap of read density for all DARs at the promoter in the CCI and Naïve groups with increased accessibility (top panels) and decreased accessibility (bottom panels). Each row represents one promoter region and the regions are aligned to the transcription start site for each gene. The color intensity represents the magnitude of chromatin accessibility. The average read density across all regions for each heatmap is shown on the right. E) Bar plot of the gene ontology analysis of the biological processes identified in promoter DARs after CCI versus Naïve.

Because the majority of chromatin accessible regions were shared in both groups, we used DiffBind to identify quantitative differences in accessibility between CCI and naïve rats. Of the 6809 promoters that contained a region of chromatin accessibility, 331 promoters were significantly more accessible and 145 were significantly less accessible in the CCI group when compared to the naïve group (Figure 2D; Supplemental table 2). GO analysis of the nearest annotated genes for these 476 differentially accessible regions (DARs) identified biological processes enriched in neuropathic pain (e.g., protein transport and assembly, cell signaling, response to stimulus, cell projection organization) (Figure 2E).

### Multi-omics analysis to identify CREs in distal intergenic regions

To identify CREs, we used ChIP-seq targeting H3K4me1. We pooled H3K4me1 enriched regions from biological duplicates of both naive and CCI rats, and identified 211,440 unique peaks. The majority of these peaks were predominantly located in intergenic and intronic regions with the remaining peaks located near or at an annotated TSS (Supplemental figure 2A). A total of 58,446 (27.6%) of these 211,440 regions overlapped with one or more accessible chromatin regions identified by our ATAC-seq, and were therefore included for further analyses (Figure 3A).

**Figure 3.**
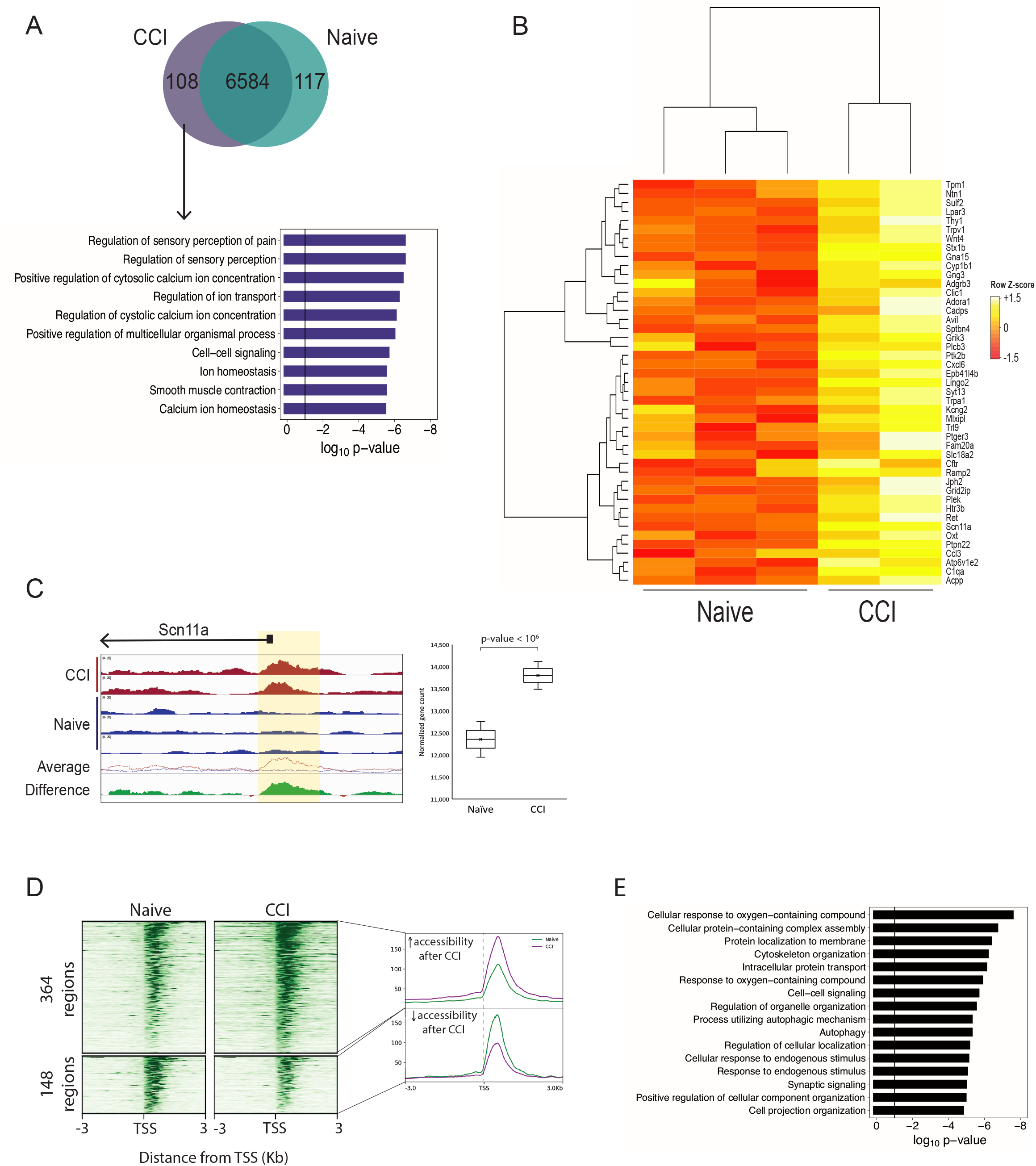
Differentially accessible regions in putative cis-regulatory regions. A) A consensus peakset of 58,446 regions showed chromatin accessibility in regions deposited with the H3K4me1 histone modification. Heatmap of the read density of each of these regions in the Naïve and CCI ATAC-seq samples and H3K4me1 ChIP-seq. Each row represents one region and the regions are aligned by its center. The color intensity represents the magnitude of read coverage. B) Volcano plot of differential accessibility of the 58, 446 accessible regions that overlap H3K4me1. Statistically significant peaks are shown in red. C) correlation heatmap of the 2145 DARs. These differential accessible regions successfully isolated regions that help us distinguish the naive from the CCI group. D) Dot plot of the significantly overrepresented motifs in intergenic DARs in the naïve DRG and after CCI. The size of the circle represents the % of DARs that contain the motif and the color indicates the q-value.

The differential analysis identified 2145 (3.67%) of the 58,446 consensus regions that showed increased or decreased accessibility between CCI and Naïve groups (Figure 3B; Supplemental table 3). PCA of only those reads contained within the 2145 DARs produced distinct clusters of the samples by group (Supplemental figure 2B) and hierarchical clustering also showed that these 2145 regions alone resulted in samples from the same treatment group clustering together (Figure 3C), which provides strong evidence that we successfully identified those genomic regions that were important in distinguishing those genomic regions affected by CCI.

Of the 2,145 DARs, 999 were located in intergenic regions (Supplemental figure 2C). Of these 999 intergenic regions, 519 (40%) had increased accessibility after CCI, and 480 had decreased accessibility. Motif analysis of the 519 DARs with increased accessibility showed enrichment for transcription factors within the bHLH/HLH, bZIP, and RUNT families (Figure 3D; Supplemental table 4). The 480 DARs with decreased accessibility after CCI were enriched in motifs from members of the high-mobility group (HMG) and the interferon regulatory factors (IRF) families (Figure 3D; Supplemental table 5). These data suggest that these transcription factors may impact susceptibility to neuron excitability through altered accessibility to their consensus DNA binding sequence.

### Changes in accessibility at CREs are associated with gene expression

To identify DARs associated with an increase or decrease in gene expression, we integrated RNA-seq data obtained from a cohort of rats that used the same experimental design (Stephens et al., 2019). We found 109 DARs with increased accessibility after CCI were located near 79 genes that were upregulated after CCI (Figure 4A). GO analysis of these 79 genes largely represented molecular functions, biological processes, and cellular compartments associated with neuronal activation and synaptic signaling (Figure 4B). A total of 39 DARs with decreased accessibility after CCI were located near 29 genes whose expression also decreased following CCI (Figure 4C). Examples of intergenic DARs associated with coordinate changes in the expression of the nearest gene are shown in Figure 4D. These findings suggest that changes in DNA accessibility at putative CREs regions following CCI can alter the expression of genes involved in nociceptive pathways.

**Figure 4.**
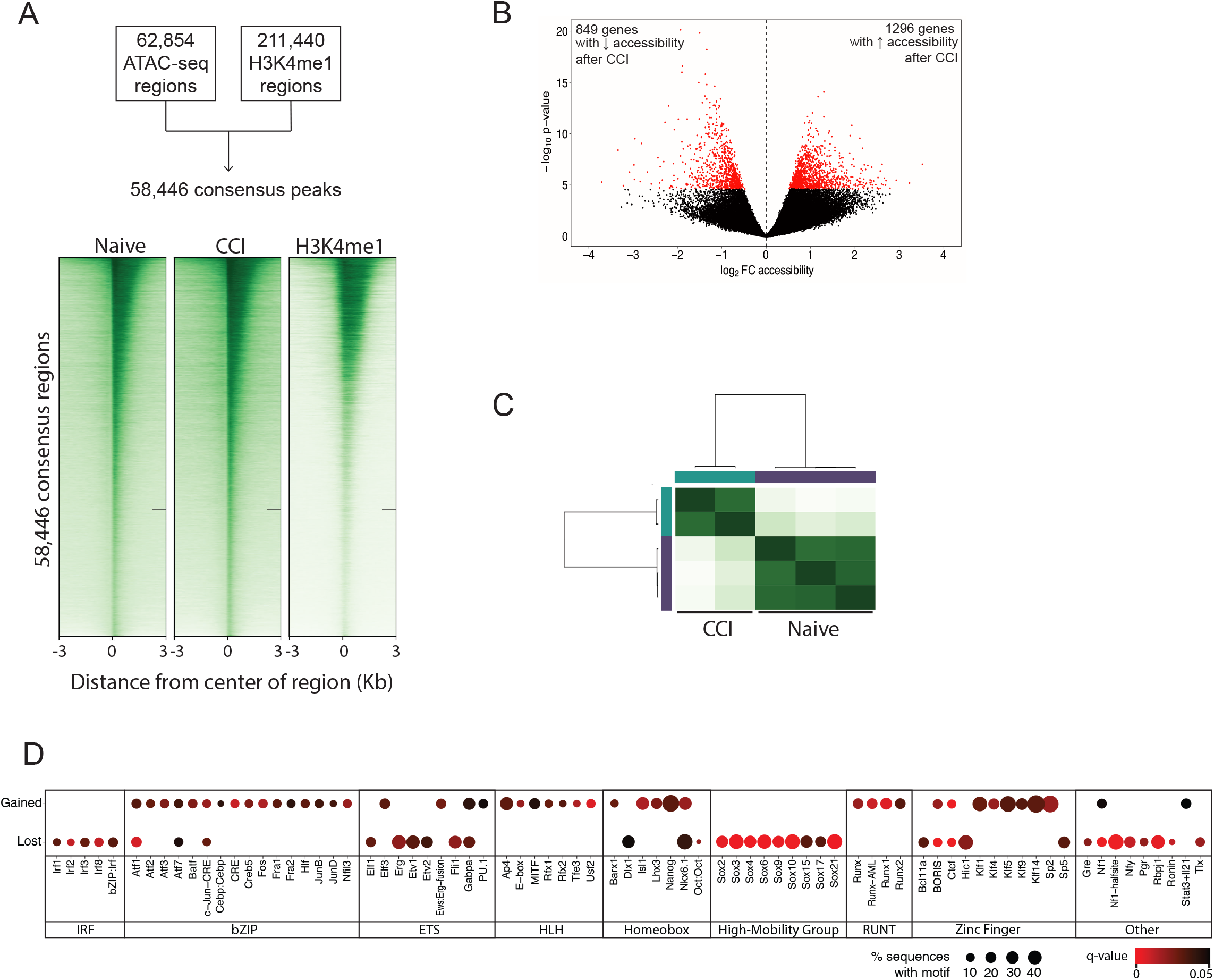
Gene expression associated with DARs. A) Scatterplot shows the FC accessibility of each region of increased accessibility and increased expression of the nearest annotated gene. Significant DARs are highlighted in red. B) Gene ontology analysis of the 79 genes with increased expression nearest to 109 regions of significantly increased chromatin accessibility. Scatterplot shows the FC accessibility of each region of increased accessibility and increased expression of the nearest gene by RNA-seq. Significant DARs are highlighted in red. Chromatin accessibility at 2 intergenic DARs (yellow highlight): 35kb upstream of Cathepsin B (left) and 31kb upstream of Cst3 (right). The individual and averaged ATAC-seq signal tracks of the normalized bigwig files for each sample as displayed from the Integrated Genomics Viewer. The “difference” track was created by subtracting the average track from the naïve group from the average track of the CCI group. Green indicates an average increase in accessibility in the CCI group compared to Naïve. The region highlighted in yellow indicates the differentially accessible region between the Naïve and CCI samples.

### Functional evaluation of CREs

To determine whether the DARs were able to alter gene expression, we selected regions associated with increased and decreased accessibility in intergenic regions. We cloned a single copy of a selected DAR into the pGL3 promoter vector which also contains the SV40 minimal promoter. Each construct was co-transfected with pGL4.74 Renilla into the 50B11 (immortalized rat nociceptor) cell line. Four constructs showed increased luciferase expression, and three showed decreased luciferase expression as compared to the empty pGL3 promoter vector (Figure 5A). These findings indicate that the DARs are associated with both enhancer (e.g., constructs A, C, E, F) and repressor (e.g., constructs B, D, G) activities, and function to increase (e.g., constructs A, C, G) or decrease gene expression (e.g., constructs B, D, E, F) in the context of physical injury.

**Figure 5.**
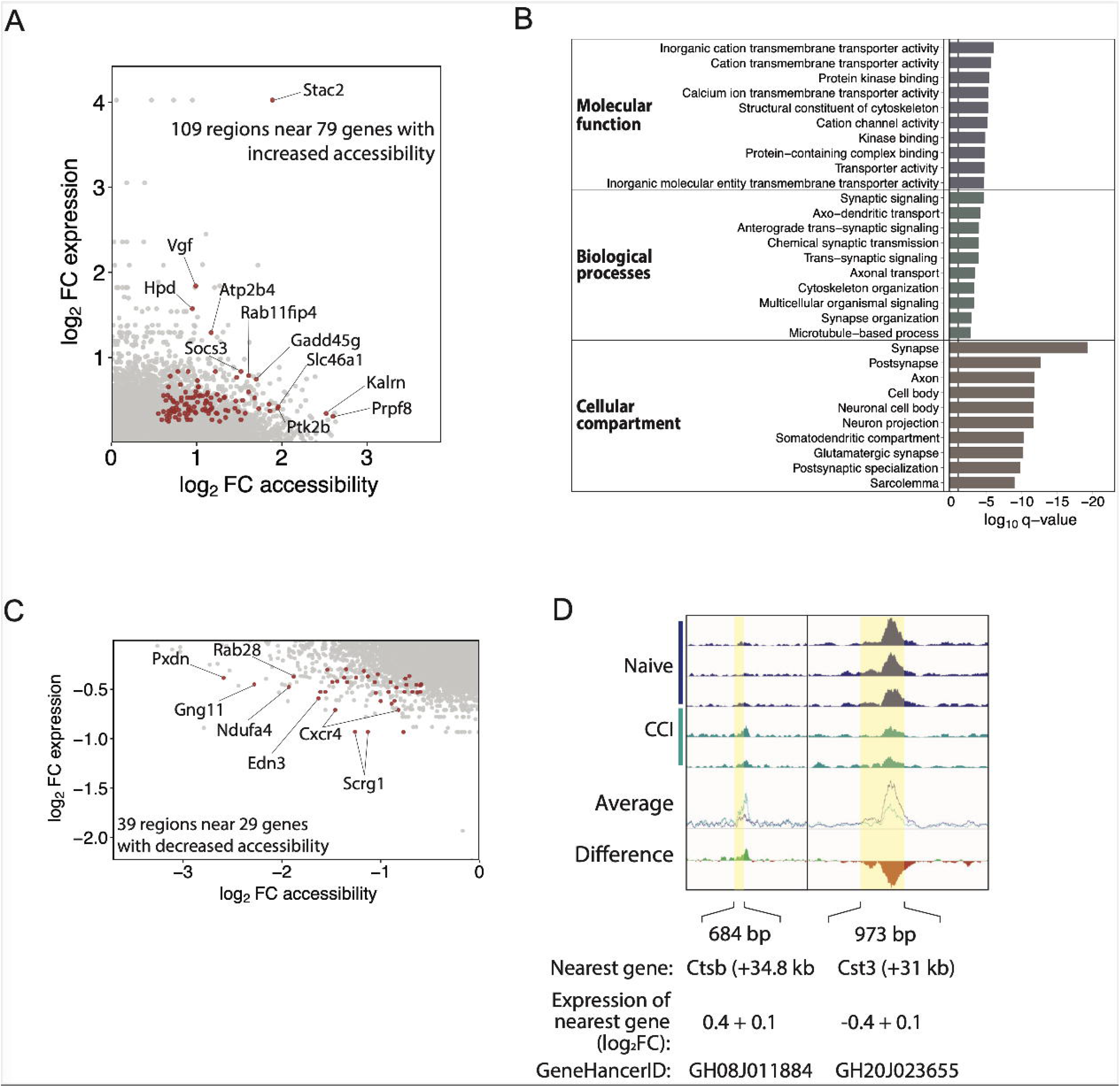
The activity of candidate DRG enhancers by luciferase reporter gene assays. Individual reporter plasmids were prepared that contained one candidate enhancer regions (A – G; Supplemental Table 8). Luciferase activity was normalized to that of the Renilla reporter and expressed as mean fold relative activity of the empty reporter ± SEM. B) Genome Viewer tracks of ATAC-seq and ChIP-seq samples chr12:12,085,353-12,088,286 of the rn6 assembly. Region highlighted in yellow corresponds to the 222bp segment cloned into the luciferase vector (Construct. G; chr12:12,086,634-12,086,856). C) 39bp segment was labeled with Cy3 and Cy5 and used to probe a human transcription factor array. Binding of CEBPG to this 39bp DNA region was verified by EMSA (D) and ChIP-PCR using primers that span chr12:12,086,663-12,086,700 (E). F) Volcano plot showing differentially expressed genes from RNAseq of 50B11 cells transduced with shRNA CEBPG knockdown or scrambled shRNA control. G) Differentially expressed genes were subjected to gene ontology analysis. REVIGO scatterplot visualizes summarized GO biological processes. Circle size is proportional to the frequency of the GO term. Color indicates the log_10_p-value.

We then determined whether the DARs showing regulatory capabilities in the luciferase assay could also bind to transcription factors by using a human transcription factor protein array (Hu et al., 2009). YBX1, HMGA1, CSDA, C/EBPγ, YBX2, CREB1, TBX3, and RHOXF1 showed significant binding to chr12:12,086,663-12,086,700 (Figure 5B-C). EMSA confirmed C/EBPγ binding to this sequence (Figure 5D). ChIP-qPCR showed significant enrichment of C/EBPγ in this DAR in 50B11 cells with a fold enrichment of 2.72 ± 0.47 over IgG (Figure 5E). Together, these studies suggested a functional role for C/EBPγ in DRG neurons in gene regulation following CCI.

### Identification of C/EBPγ-regulated genes in 50B11 cells

To identify the regulatory role of C/EBPγ in DRG neurons, we knocked down *Cebpg* expression in 50B11 cells using shRNA, and used RNA-seq to analyze gene expression changes. Western blots confirmed that *Cebpg* shRNA reduced C/EBPγ protein levels by >80%, as compared to control cells treated with scrambled shRNA (Supplemental figure 3). Compared to 50B11 cells treated with scrambled shRNA, we found 5260 differentially expressed genes (adjusted p-value < 0.01) in cells transduced with shRNA against *Cebpg* (Figure 5F). As expected, in *Cebpg* shRNA-treated cells, *Cebpg* was among the 2636 downregulated genes.REVIGO was used to summarize results from GO enrichment analysis (Figure 5G). Knockdown of *Cebpg* in 50B11 cells produced differentially expressed genes involved in various biological processes, including ion transport, transmembrane receptor signaling, regulation of trans-synaptic signaling, and regulation of cell communication.

## Discussion

By binding to accessible regulatory elements, enhancers and repressors and the chromatin remodeling proteins they recruit, provide the basis for an epigenetic pathway that prescribes specific transcriptional changes in cells when they are exposed to a subsequent stimulus (Vierbuchen et al., 2017). In this study, we determined dynamic changes of chromatin structure in DRG cells after peripheral nerve injury, which may play an important role in neuropathic pain development and maintenance. Here, we found enrichment of several AP-1 transcriptional factor family motifs in differentially accessible regions following CCI, including those bound by Atf3, Fos, Fra1, Fra2, c-Jun, JunB, and JunB. Furthermore, following CCI, we observed increased expression of At*f*3, Jun, Fos, and Jun. Consistent with prior studies, mimicking neural injury in dentate granule neurons using electroconvulsive stimulation increased the expression of AP-1 pioneer transcription factors and promoted alterations in chromatin accessibility at Fos-Jun subfamily motifs (Su et al., 2017). Interestingly, in human umbilical vein endothelial cells, ATF3 can drive increased chromatin accessibility at distal regulatory elements (Zhang et al., 2021). Together our findings suggest that increased expression of AP-1 transcription factors in DRG cells following CCI is altering the chromatin topology at key regulatory elements, and therefore, providing an avenue for prolonged neuronal plasticity following nerve injury.

### Nerve injury-induced reciprocal changes in gene expression

We highlight an example of reciprocal changes in chromatin accessibility at distinct CREs, which may act cooperatively to alter gene transcription (Figure 4D). Following CCI, a 684 bp region located 34.8 kb downstream of the gene *Ctsb* (chr15:46,281,215-46,281,898) showed a 26% increase in chromatin accessibility, and the *Ctsb* gene showed increased expression. In contrast, after CCI, chromatin accessibility was reduced at a 973 bp region located 31kb upstream of *Cst3* (chr3:143,254,480-143,255,452), and Cst3 expression increased in kind. *Ctsb* encodes Cathepsin B, a cysteine protease which promotes chronic inflammatory pain through activation of pro-caspase-1 and secretion of mature IL-1β (Sun et al., 2012), while *Cst3* encodes Cystatin C, a highly efficient cysteine protease inhibitor. The N-terminal region of Cystatin C competitively and reversibly binds to the Cathepsin B active site, thereby preventing access to potential substrates (Pavlova and Bjork, 2003). Cathepsin B and Cystatin C not only interact directly, but genetic ablation of Cystatin C increases Cathepsin B expression in, and promotes synaptic plasticity of, hippocampal neurons (Sun et al., 2008). Because Cathepsin B has been nominated as a promising therapeutic target to treat Alzheimer’s disease (Wang et al., 2012), future therapies directed towards correcting chromatin structure at respective CREs elements, like those for *Ctsb* and *Cst3*, may be able to limit potential side effects from diseases involving aberrant protein expression.

### Peripheral nerve injury may facilitate C/EBPγ mediated gene repression

Here we show that sciatic CCI is associated with an increased potential for C/EBPγ-mediated gene repression in the lumbar DRGs. C/EBPγ is a ubiquitously expressed member of the CCAAT enhancer binding (C/EBP) family, a group of bZIP transcription factors, which is involved in the regulation of major physiologic processes (e.g., control of cellular proliferation, induction of the integrated stress response, and regulation of metabolism) (Renfro et al., 2022). C/EBPγ possesses broad regulatory capacity and can activate or repress gene transcription based on its cellular context, differential chromosomal binding, and its heterodimerization with other bZIP transcription factors. In the present study, we found that CCI increased chromatin accessibility at a 222 bp region of chromosome 12 in the lumbar DRG, and that this same region repressed luciferase expression and bound C/EBPγ in a peripheral nociceptor cell line. Interestingly, Cebpg gene expression was unchanged 14 days following CCI, however, several bZIP transcription factors that are known to form heterodimers with C/EBPγ (e.g., Atf4, C/EBPβ) were upregulated. Future functional studies may help determine whether these or other transcription factors heterodimerize with C/EBPγ at regions of increased chromatin accessibility to alter the neuronal response to injury.

Given the context-dependent activity of C/EBPγ, its functional roles in developing and mature nervous system cells are poorly understood. To broadly identify genes that C/EBPγ may be regulating following nerve injury, we knocked down *Cebpg* in a rat nociceptor cell line and found differentially expressed genes associated with a variety of pain-related biological processes including carboxylic acid transport, trans-synaptic signaling, signaling receptor activity, and ion homeostasis. Our data supports existing literature regarding involvement of C/EBPγ in mediating the development of chronic pain (Renfro et al., 2022). In the sciatic crush nerve injury model, *Cebpg* expression was significantly upregulated in the DRG at approximately 1-day post-injury before returning to baseline levels after 3 days (Lopez de Heredia and Magoulas, 2013). A second study also implicates *Cebpg* expression in the development of mechanical hypersensitivity following spared nerve injury (Mamet et al., 2014). Prior to sciatic nerve injury (SNI), rats were treated with AYX1, a DNA decoy that targets Egr1. *Cebpg*, which is regulated by Egr1, was significantly downregulated in the DRG and spinal cord of AYX1-treated rats, which also showed reduced mechanical hypersensitivity following a perhipheral nerve injury. Further research is needed to better understand the impact of C/EBPγ-mediated gene repression in the DRG following CCI and during the development and maintenance of pain hypersensitivity.

In conclusion, we used ATAC-seq and ChIP-seq to identify accessible chromatin regions associated with regulatory regions in the genome of rat DRGs. Our multiomic assessment of nerve-injured and naïve DRG provides novel insights into the role of chromatin structure at regulatory elements and activities of transcription factors that together may facilitate neuronal excitability and nerve regeneration. Our improved understanding of how epigenetic perturbations alter transcription in response to nerve injury, and their relationship to chronic pain will facilitate the development of novel classes of analgesics that can target these and other similar mechanisms upstream of transcription.

## Acknowledgements

This study was supported by grants from the National Institutes of Health (Bethesda, Maryland, USA) F32NR015728 (KES), KL2 TR003108 (KES), P20GM121293 (KES), NS070814 (YG), NS110598 (YG), NS117761 (YG), R01GM118760 (SDT), R01CA221306, and the National Science Foundation (Alexandria, Virginia, USA) 132452 (SDT), the Arkansas Children’s Research Institute (KES), the Arkansas Breast Cancer Research Program (KES) as well as a seed grant from the Johns Hopkins Blaustein Pain Research Fund (SDT). We thank Rakel Tryggvadóttir for her technical assistance. All authors report no conflicts of interest.

## Author contributions

Conceptualization, K.E.S., Y.G., and S.D.T.; Methodology, K.E.S., Y.G., and S.D.T.; Validation, C.M., Z.R. and D.A.V.; Formal Analysis, K.E.S., W.Z., C.Z., and H.J.; Investigation, K.E.S., C.M., D.A.V., B.E.W., Z.R.; Resources, K.E.S., H.K., H.Z., Y.G. and S.D.T.; Writing – Original Draft, K.E.S.; Writing – Review & Editing, K.E.S, Y.G., S.D.T, H.Z., W.Z.; Visualization, K.E.S.; Supervision, K.E.S., H.Z., Y.G. and S.D.T.; Funding Acquisition, K.E.S., Y.G., and S.D.T.

## Data and Code Availability

Raw and processed sequencing data for all ATAC-seq data and the CHIP-seq data have been deposited in the NCBI GEO database under accession #GSE210321. The RNA-seq data files for the naïve and CCI groups were previously published and are available under accession #GEO100122 (Stephens et al., 2019).

## Figure legends

**Supplemental figure 1**. A) Principal component analysis plot of ATAC-seq biological replicates for the naïve (aqua; n=3) and CCI (purple; n=2) groups. The percent variance that the first and second principal components explain are included in the axis labels. B) The total number of individual ATAC-seq peaks identified in each sample is provided in the table on the left. The consensus peakset consists of 62,854 of these regions where peaks were called in at least 40% of samples (i.e., minimum 3 of 5). The table on the right indicates the number of regions in the consensus peakset that were found only in the CCI (i.e., peak present in 2 CCI samples and 0 naïve samples), naïve (i.e., peaks present in 2 or 3 of the naïve samples and 0 CCI samples), or in both groups (i.e., peaks present in at least 1 sample from each group). C) Distance of accessible regions from nearest annotated gene. The distribution of accessible regions shows their concentration in cis-regulatory regions versus uniform/random distribution along the genome. D) Classification of accessible regions by genomic feature for regions present in only CCI, only Naive, or both CCI and naive. The colored bars represent the percentage of the total number of regions for specific features. The black bars indicate the total number of regions.

**Supplemental figure 2**. A) H3K4me1 peaks by feature. B) Principal component analysis of the 2145 DARs. C) Genomic features associated with each of the 2145 DARs that identify features with increased (i.e., gained) accessibility or decreased (i.e., lost) accessibility after CCI as compared with Naïve.

**Supplemental figure 3**. Western blot of Cebpg shRNA-mediated knockdown following transfection of pGFP-C-shLenti in 50B11 cells. A) Representative band expression of the load control β-actin for normalization of protein loading (Top) and a comparison of the Control shRNA scrambled sequence (ShS) with three different shRNA constructs (ShA, ShC, and ShD) used to achieve sufficient knockdown (>70%) (Bottom). B) Average normalized band density. Constructs (ShA, ShC, and ShD) were statistically compared to the control scrambled sequence (ShS) using Analysis of Variance followed by a Dunnett’s test for multiple comparisons. (Significant difference is based on p< 0.05, *** represents a p-value that is less than 0.0005, Error bars represent Standard Deviation). C) Original images of western blots.

## Tables

Supplemental table 1 – ATAC-seq/ChIP-seq/RNA-seq sequencing metrics

Supplemental table 2 – Promoter regions with increased and decreased chromatin accessibility

Supplemental table 3 – H3K4me1 regions with increased and decreased accessibility after nerve injury.

Supplemental table 4 – List of transcription factor motifs enriched in genomic regions that increased in accessibility following CCI

Supplemental table 5 –List of transcription factor motifs enriched in genomic regions that decreased in accessibility following CCI

Supplemental table 6 – Locations of DARs for luciferase assay with primers

## Methods

### Animals

Adult female Sprague-Dawley rats (12-16 weeks old) (Harlan Bioproducts for Science, Indianapolis, IN) were housed 2-3 per cage in centralized animal care facilities with 12-hour light/dark cycle. Animals were allowed to acclimate for a minimum of 48 hours prior to any procedures and given *ad libitum* access to food and water. All procedures involving animals were reviewed and approved by the Johns Hopkins Animal Care and Use Committee and are performed in accordance with the NIH Guide for the Care and Use of Laboratory Animals.

### CCI of the sciatic nerve

CCI surgery to the sciatic nerve was performed on all rats as previously described (Bennett and Xie, 1988). Under 2-3% isoflurane, a small incision was made at the level of the mid-thigh. The sciatic nerve was exposed by blunt dissection through the biceps femoris. The nerve trunk proximal to the distal branching point was loosely ligated with four 4-O silk sutures placed approximately 0.5mm apart until the epineuria was slightly compressed and minor twitching of the relevant muscles was observed. The muscle layer was closed with 4-O silk suture and the wound was closed with metal clips. All surgical procedures were performed by the same individual to avoid variation in technique. Hypersensitivity of the hind paws was verified by von Frey monofilaments (Chaplan et al., 1994) on day 14 post-injury.

### Experimental design

Rats were assigned randomly to either receive CCI surgery or no procedure (i.e., naive control) (Figure 1A). On postoperative day 14 rats were euthanized by overdose of isoflurane and decapitation after which the ipsilateral L4-L6 DRGs were quickly dissected. For ChIP-seq and RNA-seq, DRGs were immediately submerged in liquid nitrogen and stored at -80°C until processing.

### ChIP-seq library preparation

Chromatin immunoprecipitation (ChIP) was used to identify sequences enriched with H3K4me1 in the rat DRG. The ipsilateral L4-L6 DRGs harvested from 3 animals were pooled and cross-linked in 1% formaldehyde for 10 minutes at room temperature. The DRGs were washed in 1X PBS and subjected to dounce homogenization in a lysis buffer (0.32M sucrose, 5mM CaCl_2_, 3mM Mg(Acetate)_2_, 0.1mM EDTA, 10mM Tris-HCl, pH 8.0, 1mM DTT, 0.1% Triton X-100). The homogenate was centrifuged for 10 minutes at 10,00xg at 4°C to pellet the nuclei. Nuclei were resuspended in a nuclei lysis buffer (25mM Tris-HCl, pH 8.0, 5mM EDTA, 1% SDS). The chromatin was sheared using the Bioruptor sonicator (Diagenode; Liege, Belgium) with high output and 35 cycles of 30 seconds on/30 seconds off to produce DNA fragments with lengths between 200-600 base pairs. Sheared chromatin was then diluted in RIPA buffer (10mM Tris-HCl, pH 8.0, 1mM EDTA, 1% Triton X-100, 0.1% SDS, 0.1% Na deoxycholate, 100mM NaCl) and incubated with anti-H3K4me1 antibody (ab8895; abcam; Cambridge, MA) attached to protein G Dynabeads (Invitrogen). The chromatin-bead preparation was incubated at 4°C for 2 hours. An aliquot of sheared chromatin was taken as the input sample (i.e., pre-precipitation control). Following incubation, each immunoprecipitation reaction was washed 3 times with low salt wash buffer (20mM Tris-HCl, pH 8.0, 2mM EDTA, 0.1% SDS, 1% Triton X-100, 150mM NaCl) and once with high salt wash buffer (20mM Tris-HCl, pH 8.0, 2mM EDTA, 0.1% SDS, 1% Triton X-100, 500mM NaCl). The DNA-histone complexes were eluted from the Dynabeads (1% SDS, 100mM NaHCO_3_). Cross-links between the DNA fragments and histones were reversed and the DNA fragments were recovered using the ChIP DNA Clean & Concentrator Kit (Zymo, Irvine, CA). Two biological replicates were performed for each group. The input sample and enriched DNA samples obtained from the ChIP assays were used for library construction using the TruSeq ChIP Sample Prep Kit (Illumina; San Diego, CA) according to manufacturer’s instructions. Libraries were quantified using the KAPA qPCR quantification kit (KAPA Biosystems, Wilmington, MA) and sequenced on an Illumina HiSeq 2500 producing single end 50 base pair reads. All pre-immunoprecipitation buffers contained protease inhibitors (1mM Benzamidine, 1mM PMSF, 5mM Na Butyrate).

### ATAC-seq library preparation

Immediately following dissection, the ipsilateral L4-L6 DRGs from one rat were transferred directly to cold lysis buffer (0.32M sucrose, 5mM CaCl_2_, 3mM Mg(Acetate), 10mM Tris-HCl, pH 8.0, 0.1% Triton X-100, 1mM DTT, 5mM Na Butyrate, 1mM PMSF). Nuclei were isolated through dounce homogenization of the tissue in lysis buffer followed by ultracentrifugation through a sucrose cushion (1.8M sucrose, 3mM Mg(Acetate), 1mM DTT, 10mM Tris-HCl, pH 8.0, 5mM Na Butyrate, 1mM PMSF) at 139,800 x g at 4° C for 2 hours to remove mitochondrial DNA. The nuclei were resuspended in 1X PBS and counted 3 times using a Neubauer chamber. Tagmentation by Tn5 was performed using reagents from the Nextera DNA Sample Preparation Kit (FC-121-1030, Illumina; San Diego, CA) as previously described (Buenrostro et al., 2013). Each 50ul reaction contained 50,000 nuclei, 25ul 2X Tagmentation Buffer, and 2.5ul Tn5 enzyme and incubated at 37° C for 30 minutes. Tagmented DNA was immediately purified using the Clean and Concentrate-5 Kit (Zymo, Irvine, CA) and eluted in 10ul elution buffer. Tagmented DNA fragments were amplified using Nextera Index adapters, PCR primer cocktail, NPM PCR master mix and 10 cycles of PCR. Each library was purified using Agencourt AMPure XP beads (Beckam Coulter; Atlanta, GA). The fragment distribution of each library was assessed using the High Sensitivity DNA Kit on an Agilent 2100 Bioanalyzer (Agilent Technologies; Palo Alto, CA). Libraries were quantified prior to sequencing using the Qubit DNA HS kit (ThermoScientific, Waltham, MA) and normalized to 2nM and pooled in equimolar concentrations. Libraries were sequenced using paired-end, dual-index sequencing on an Illumina HiSeq 2500 (Illumina, San Diego, CA) which produced 50bp reads. ATAC-seq was performed on 3 biological replicates for the CCI group and 3 biological replicates for the naive group.

### RNA-seq dataset

We performed RNA-seq from DRGs from a similar cohort of adult, naïve female Sprague-Dawley rats or after CCI and validated this data using qPCR in biological replicates. Details regarding sample acquisition, RNA library preparation, and RNA-seq data processing have been published (Stephens et al., 2019). RNA-seq data are available under accession #GEO100122.

### 50B11 cell culture

The 50B11 cells were a gift from Ahmet Höke of the Johns Hopkins University, Department of Neurosurgery. Cells were maintained in Neurobasal medium (ThermoScientific, Waltham, MA) supplemented with 2% B27 (Sigma; St. Louis, MO), 550µM glutamine (Sigma; St. Louis, MO), 12mM glucose, and 10% fetal bovine serum and incubated at 37°C in a humidified environment containing 5% CO_2_. Cells with low passage numbers (i.e. <20) were used for all experiments.

### Cloning

Luciferase reporter constructs were generated by cloning a candidate enhancer region into the pGL3 promoter vector (Promega; Madison, WI). Each region was inserted using standard restriction enzyme-based cloning techniques. The regions were obtained by PCR of rat genomic DNA. The 5’ end of the primers were modified to contain BglII (forward primer) and MluI (reverse primer) restriction sites (Supplemental table 6). PCR was performed using the Pfu Turbo polymerase (Agilent Technologies; Palo Alto, CA) and touchdown thermocycling. The PCR products were digested and ligated into the BglII (AGATCT) and MluI (ACGCGT) restriction enzyme sites of the pGL3-Promoter luciferase vector (Promega; Madison, WI). The ligated products were transformed into chemically competent DH5α cells using ampicillin (100mcg/ml) to select for the recombinant plasmid-positive colonies. All constructs were verified by diagnostic restriction enzyme digestion and Sanger sequencing.

### Transfection and luciferase assays

50B11 cells were seeded at 10,000 cells/well in 48 well plates in 250ul of complete media and grown to 60-80% confluence. Cells were then transfected with each reporter construct (450ng) and 50ng pGL4.74 Renilla luciferase expression vector (Promega; Madison, WI) using ViaFect Transfection Reagent (Promega; Madison. WI) in 25ul Opti-MEM (ThermoScientific, Waltham, MA) with a 4:1 ratio in 250µl complete medium. The transfection efficiency of 50B11 cells was evaluated by transfecting cells with EGFP-N1 (Clontech; Mountain View, CA) in parallel reactions. At 48 hours post-transfection, Firefly and Renilla luciferase were measured using the Dual-Glo Luciferase reporter assay system (Promega). The amount of Firefly luciferase normalized to the Renilla luciferase and expressed as the relative fold difference of the empty pGL3 promoter vector. Each enhancer construct was tested in quadruplicate.

### Transcription factor protein microarrays

Human transcription factors were purified from yeast as GST fusion and arrayed on FAST slides in duplicate as previously described (Hu et al., 2013, Hu et al., 2009). The microarrays were probed with a 39-nucleotide sequence from differentially accessible regions used in the luciferase assays. Sixty base oligonucleotides were synthesized by IDT and contained this 39-nucleotide sequence followed by the reverse T7 sequence. The DNA probes were converted to double-stranded DNA with either Cy3- or Cy5-labeled T7 primer as the 5’-end. The labeled double-stranded T7 sequence was chosen as a negative control for T7-specific binding. Each sequence was tested in duplicate arrays with alternating fluorescent labels. The slides were then washed and scanned with a GenePix 4000B scanner (Molecular Devices, Sunnyvale, CA) and the binding signals were acquired using the GenePix 6.0 software. GenePix 6.0 was used to align the spot-calling grid and record the foreground and background intensities for every protein spot. The raw binding intensity for each probe was defined as F_ij_/B_ij_, where F_ij_ and B_ij_ are the median values of foreground and background signals of the probes at site (i,j) on the microarray, respectively. We normalized the raw signal of each probe based on the median value of raw signals of its neighboring probes. The Z-score of each binding assay was calculated by Z_i,j_ = (R’_I,j_ – Ñ)/std(N), where R’_I,j_ is the locally normalized intensity of probe (I,j) on the microarray, Ñ and std(N) are mean value and standard deviation, respectively, of noise distribution on the microarray. Since each protein is printed in duplicate on the microarray and each binding assay was performed in duplicate, a protein was identified as a positive hit only when all of its 4 spots produced a Z-score > 3.0.

### EMSA

The binding reaction was carried out with 100 fmol of biotinylated dsDNA probe and 1 pmol of purified C/EBPγ protein in 20µl of binding buffer as previously described (Hu et al., 2013, Hu et al., 2009). 10-fold of unlabeled T7 was added in the competition assay. The C/EBPγ expression clone used in the EMSA was verified by DNA sequencing.

### ChIP-qPCR in 50B11 cells

ChIP-qPCR was used quantify Cebpg binding to the putative enhancer element (rn6:chr12:12,085,353-12,088,386) in 50B11 cells. Five million 50B11 cells were cross-linked in 1% formaldehyde for 10 minutes at room temperature. A final concentration of 0.125M glycine was added to quench the cross-linking reaction. The cells were washed in chilled 1X PBS and subjected to dounce homogenization in lysis buffer (5mM HEPES, 85mM KCl, 1% IGEPAL CA-630 in 1X PBS, 1X Roche Complete protease inhibitors, 5mM Na Butyrate). The homogenate was centrifuged for 10 minutes at 10,00xg at 4°C to pellet the nuclei. Nuclei were resuspended in nuclei lysis buffer (5mM Tris-HCl, pH 8.0, 10mM EDTA, 1% SDS in 1X PBS, 1X Roche Complete protease inhibitors, 5mM Na Butyrate) and incubated at room temperature for 10 minutes. The chromatin was then sheared using a Qsonica Q800R2 (Qsonica, Newtown, CT) at 75% power for 15 cycles of 30 seconds on/30 seconds off to produce DNA fragments with lengths between 200-600 base pairs. Chromatin was then stored at -80°C until immunoprecipitation. Sheared chromatin was incubated with anti-CEBPG or anti-rabbit IgG control antibody attached to protein A Dynabeads (Invitrogen) and incubated overnight at 4°C. An aliquot of sheared chromatin was taken as the input sample (i.e., pre-precipitation control). Following incubation, each immunoprecipitation reaction was washed 3 times with low salt wash buffer (50mM Tris-HCl, pH 7.4, 10mM EDTA, 0.1% SDS, 1% Triton X-100, 150mM NaCl, 1mM PMSF), five times with high salt wash buffer (50mM Tris, pH 7.4, 500mM NaCl, 1% Triton-X-100, 0.1% SDS, 10mM EDTA, 1mM PMSF), and once with 250mM LiCl, 50mM Tris, pH 7.4. The DNA-histone complexes were eluted from the Dynabeads (1% SDS, 100mM NaHCO_3_) at 65°C. Cross-links between the DNA fragments and histones were reversed. and the DNA fragments were recovered using the ChIP DNA Clean & Concentrator Kit (Zymo, Irvine, CA) and quantified by Qubit. Each 10 µl qPCR reaction consisted of 2X SSoAdvanced Universal SYBR green qPCR Master Mix (Bio-Rad, Hercules, CA), 200 nM each forward and reverse primer, and 3.2 µl ChIP DNA. PCR of each target was performed in triplicate using the CFX384 Touch Real-Time PCR Detection System (Bio-Rad, Hercules, CA) with the following thermocycling conditions: initial denaturation at 98°C for 3 minutes followed by 40 cycles of 98°C for 15 seconds and 60°C for 22 seconds. Nuclease-free water was included as the no-template control. Ct values were used to calculate the percent input/fold enrichment above the IgG/NTC controls. Primers used to amplify the 5’ region of Cebpg were CCT TGA GGG TTC TTC GGC TG (forward) and CTG TGG TGT GCT CGA GTG AT (reverse).

### Generation of CEBPG knockdown in 50B11 cells

Stable knockdown of Cebpg in 50B11 cells was done by transfecting 50B11 cells with pGFP-C-shLenti carrying shRNA against Cebpg (1µg/ml; TL709448; Origene, Rockville, MD) following manufacturer’s recommendations. A 29-mer scrambled shRNA cassette in the pGFP-C-shLenti vector (TR30021; Origene) was used as the off-target control. Transduced cells then underwent puromycin selection and visible confirmation for GFP. Cebpg knockdown efficiency compared to control was verified by Western blot (Supplemental figure 3).

### Western blot

Whole-cell lysates (50µg/sample) were run on 12% SDS-PAGE and transferred to a nitrocellulose membrane. The membrane was blocked for 1 hour with 5% blocking buffer (Bio Rad, Cat# 1706404) and incubated overnight at 4°C with primary rabbit polyclonal anti-CEBPG antibody (1:1000; MyBioSource, Cat# MBS8241686) and anti-β-actin antibody (1:1000; Cell Signaling, Cat# 4970S). Targets were detected after incubation with a goat anti-rabbit secondary antibody horseradish peroxidase conjugated (1:10,000; Cell Signaling, Cat# 7074P2) for 1 hour and visualized with the chemiluminescence reagent (Immobilon Forte Western HRP Substrate; Millipore, Cat# WBLOF0100). Images of the membrane were captured by the ImageQuant LAS 4000 (GE Healthcare life Sciences).

### RNA-seq

Total RNA was extracted from transgenic Cebpg knockdown and control 50B11 cell lines using trizol-chloroform. The RNA Clean and Concentrate-5 kit was used to clean up RNA from the aqueous phase with on column DNAseI digestion. RNA concentration was measured using the NanoDrop One Spectrophotometer (Thermo Fisher Scientific, Waltham, MA) and RNA integrity was assessed using RNA Nano Eukaryote chips in an Agilent 2100 Bioanalyzer (Agilent Technologies, Palo Alto, CA). One microgram of total RNA was used to construct sequencing libraries as previously described (Stephens et al., 2019, Stephens et al., 2021). Strand-specific RNA libraries were prepared using the NEBNext Ultra II Directional RNA Library Prep Kit for Illumina with NEBNEXT poly(A) mRNA Isolation Module (New England Biolabs) according to manufacturer recommendations. Samples were barcoded using the recommended NEBNext Multiplex Oligos (New England Biolabs). The size range and quality of libraries were verified on the Agilent 2100 Bioanalyzer (Agilent Technologies, Palo Alto, CA). RNA-seq libraries were quantified by qPCR using the KAPA library quantification kit (KAPA Biosystems, Wilmington, MA). Each library was normalized to 2nM and pooled in equimolar concentrations. Single-end sequencing (i.e, 1×75bp) was performed in a single run on an Illumina NextSeq 500 (Illumina, San Diego, CA). Three independent experimental replicates were run for the Cebpg shRNA-mediated knockdown and shRNA control transgenic lines for a total of 6 libraries.

### Data analysis

#### ChIP-seq

Raw fastq files were aligned to rat RN6 genome using Bowtie2 (Langmead and Salzberg, 2012). Duplicated reads were removed using Picard tools (2019). Peak calling was performed with MACS2 (Zhang et al., 2008). Regions of H3K4me1 enrichment were identified by MACS2 for the CCI biological duplicates and separately for the Naive biological duplicates using input samples as controls and the following settings —keep-dup all -B —SPRM — nomodel —broad. All regions found to be enriched in either group were merged into a single list and overlapping regions reduced to be expressed as a single region.

#### ATAC-seq

Paired-end reads were trimmed using Trimmomatic (Bolger et al., 2014) to remove adaptors. The trimmed reads were then aligned to rat genome rn6 using Bowtie2 (Langmead and Salzberg, 2012) with the following parameters -X2000 --no-mixed --no-discordant. Reads with a mapping quality score less than 10 were removed using SAMtools (Li et al., 2009) and duplicated reads were removed using the MarkDuplicates function in Picard. The genomic coordinates for each read were then shifted 4 bases (positive strand) or 5 bases (negative strand) 5’ relative to the reference genome to adjust each read for the Tn5 binding footprint (Buenrostro et al., 2013). Each read was then trimmed to produce a single base located at the 5’ end of each read. Each single base read was extended 75 bases in each direction so that the Tn5 insertion site was located at the center of a 150-base read. These shifted reads were provided as input for peak calling with MACS2 using the following parameters: -nomodel - extsize 150 -B -keep-dup all -call-summits. The read density was calculated in 300 nucleotide bins across the genome for each sample.

#### Visualization

Tracks for each sample were created for visualization in IGV. The number of slopped insertion sites for each sample was downsampled to 30 million and converted to bigWig files using ucsctools. To create an aggregated track for each treatment group, all slopped insertion sites were concatenated into a single file and downsampled to 125 million sites for each group. These files were then converted into bigWig format. In IGV, these aggregated tracks were subtracted to create an additional track to visualize regions of increased and decreased DNA accessibility.

#### Identification of differentially accessible regions

Only regions where both H3K4me1 enrichment and accessibility in one or more samples by ATAC-seq were used to identify regions of differential accessibility. To obtain a consensus list of candidate regions, ATAC-seq peaks from each sample were subsetted by the regions of H3K4me1 enrichment. These regions were evaluated for differential accessibility using DiffBind (Ross-Innes et al., 2012). To increase the stringency of the analysis, an ATAC-seq peak must have been present in a minimum of 2 samples per group. Therefore, the ATAC-seq peak was required to be present in both CCI samples and 2 of the 3 naive samples for inclusion into analysis. Because the contribution of any single region to the CCI phenotype is predicted to be relatively small, we conducted the differential analysis using a permissive significance threshold of p< 0.05 to avoid missing potentially important contributors. The results were annotated using HOMER according to default parameters and merged with the RNA-seq data. Regions that were associated with increased or decreased gene expression were identified. Gene lists were used as input for GO analysis of biological function using the ToppGeneSuite (Chen et al., 2009). Motif analysis was conducted using the findMotifsGenome.pl command in HOMER and the rn6 genome with the following parameters: -size 200 -l 6,8,10,12.

#### RNA-seq of 50B11 transgenic lines

Sequencing reads were aligned to annotated RefSeq genes in the rat reference genome (rn6) using HISAT2 (Kim et al., 2015) and filtered to remove ribosomal RNA. A gene count matrix that contained raw transcript counts for each annotated gene was generated using the *featureCounts* function of the Subread package in R (Liao et al., 2014) against the Ensemble rn6 transcriptome. This count matrix was then filtered for low count genes so that only those genes with >0 reads across all samples were retained. We relied on the automatic and independent filtering used by DESeq2 to determine the most appropriate threshold for removing genes with low counts (Love et al., 2014). To identify genes that were differentially regulated with Cepbg shRNA-mediated knockdown, raw transcript counts were normalized, log_2_ transformed, and analyzed using the default procedures in DESeq2 (Love et al., 2014). Adjusted p-values were corrected using the Benjamini-Hochberg method. An adjusted p-value <0.05 and an absolute log_2_ fold change > 0.5 were used to define differentially expressed genes between knockdown and control. REVIGO was used to reduce and visualize GO enrichment data (Supek et al., 2011).

